# Male mice prefer to live on their own

**DOI:** 10.1101/2025.05.02.651815

**Authors:** Jennifer Davies, Megan G Jackson, Justyna K Hinchcliffe, Michael Mendl, Emma SJ Robinson

**Affiliations:** School of Physiology, Pharmacology & Neuroscience, University of Bristol, Biomedical Sciences Building, University Walk, Bristol, BS8 1TD, UK; School of Veterinary Sciences, University of Bristol, Langford House, Langford, BS40 5DU, UK

**Keywords:** Animal welfare, affective state, ultrasonic vocalisations, reward learning, 3Rs

## Abstract

Around the world, guidance for the management of laboratory mice recommends social housing (1-3). However, fighting in male mice represents a major animal welfare and scientific issue. Despite the risks of stress and injury and associated welfare costs of incage aggression, we currently prioritise group housing. In this study we first validated a trans-species behavioural biomarker of affective state for mice using a chronic corticosterone model of depression. We next used the same methods to quantify affective state in male mice, group or singly housed in conventional versus individually ventilated cages (IVCs). The results revealed group housed male mice are in a negative affective state similar to that induced by chronic corticosterone treatment. IVC housing was also associated with a more negative affective state. These studies provide an objective assessment of the welfare costs of different housing conditions for male mice.

---

Standard housing for male laboratory mice is associated with aggression and fighting which can result in serious injury or even death (4-6). Animals may also experience chronic social stress which does not result in overt injury but can still negatively impact animal welfare (7). The severity of these interactions and associated chronic stress, pain and suffering has the potential to lead to cumulative suffering which, if they were to occur as part of an experimental procedure, could reach severity limits of moderate or even severe. Stress may also increase variability of biological data reducing effect sizes and reproducibility (8, 9). Despite a literature on social behaviour in mice, there is a lack of clear recommendations for how to best manage male laboratory mice. The obvious way to avoid aggression is to individually house male mice but the consensus has been that mice are gregarious and social housing should be used where possible in reduce negative welfare effects arising from social isolation (1-3).

Given the wide acceptance that affective states are the key determinants of animal welfare, the only way to assess the welfare costs of group versus individual housing in male mice is to quantify these states (10, 11). Methods such as measures of stress hormones or injuries lack sensitivity and specificity (12). In 2004, Mendl and colleagues published the first report showing that animals in a putative negative affective state exhibit pessimistic decision-making in a behavioural task (13). This work set a framework for the application of these behavioural approaches as objective methods to quantify affective states in non-human species (14). One trans-species cognitive bias characterised in humans and rats is a negative bias in reward learning (15). In the rat reward learning assay, subjects learn that one cue (a digging substrate) predicts finding a higher value reward and then use this information in a preference test to bias their responding i.e. they develop a reward-induced positive bias. However, animals in a putative negative affective state fail to show this (16, 17).

## Development and validation of a reward bias assay for mice

The rat reward learning task involves a bowl digging protocol which is difficult to perform in mice due to their affective response to human handling. We therefore first developed a semi-automated reward learning task for mice, the reward bias assay. The task quantified reward learning over time with animals presented with pairs of cues (textured floors) predicting high value versus no reward or low value versus no reward. The cues were presented using a T-maze apparatus (illustrated in Figure 1A) and with the two pairs of cues presented in a pseudorandom spatial order within each session. Animals were pre-trained over 5 days to confirm that they could discriminate between texture vs plain floor types, before being tested for discrimination learning of different texture cues predicting high versus low reward over 10 sessions of 20 trials per session (Figure 1B). Accuracy for discrimination between each of the pairs of cues was calculated, plotted against time and analysed using a RM 2-way ANOVA or mixed effects analysis with time (session) and reward value (reward) as factors. To validate the method, single housed male C57/Bl (n = 18, 8 weeks old) mice were treated with corticosterone or vehicle (25ug/ml corticosterone-HBC in the drinking water) for 14 days to induce a depression-like state (18). Treatment was continued while all animals were tested in the reward bias assay. As summarised in figure 1C and 1D, the animals treated with chronic corticosterone exhibited reduced reward-induced biases, failing to develop a higher accuracy for the high reward association. Over the 10 sessions, control animals showed a trend towards learning over time (F_(3.892, 31.14)_ = 2.667, p = 0.052) for both rewarded cues, and exhibited higher accuracy over time for the high reward association, and a main effect of reward value (F_(1, 8)_ = 16.61, p = 0.0036). In contrast, mice treated with chronic corticosterone showed no evidence of learning over time (P > 0.05) or effect of reward value (p > 0.05). Comparison of the reward-induced bias on session 10 found a reduction in bias score for the corticosterone-treated animals compared to controls (U _(54, 24)_ = 3, p = 0.0216, Mann Whitney Test). The impairment observed in the chronic corticosterone treated mice was similar to that observed in rat models of depression(16, 17, 19-21).

**Fig. 1.**
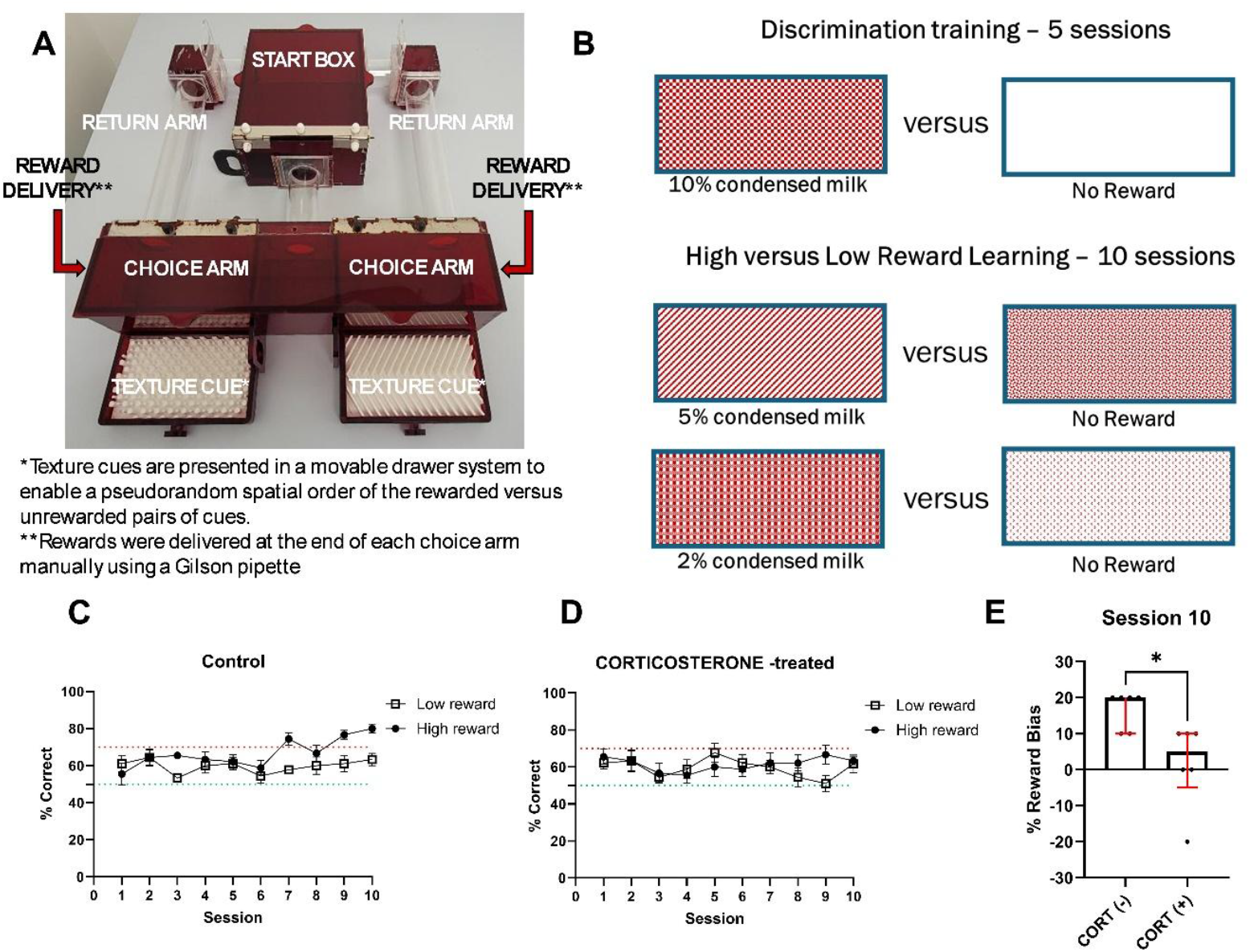
Mice treated with chronic corticosterone to induce a negative affective state show impaired reward learning in the reward bias assay. A. Photograph of the Complex Cue T-maze (CCTM) apparatus (GB Patent Application No: 2306529.5). Each arm of the CCTM has two interchangeable texture floors which can be presented in a pseudorandom spatial order. B. Illustration of the pairs of cues used for the reward bias assay. Pairs of cues were consistent throughout the study with each pair including a rewarded (high or low value reward) and unrewarded texture cue. Cues were counter-balanced across subjects and reward value but remained consistent for each subject across all sessions. Validation data is illustrated in C-E. Mice underwent chronic corticosterone or vehicle treatment in the drinking water for a period of two weeks (25 µg/ml). They were then tested in the reward bias assay over 10 sessions. **C&D** shows learning accuracy across conditions for the high versus low reward. **E** corticosterone treatment reduced reward bias on session 10. CORT=corticosterone. *P < 0.05. Bars are mean ± SEM (C and D) or median and interquartile with data points overlaid (E, n = 9 per group, n = 6 on session 10).

## Male mice housed in groups and IVC caging systems are in a more negative affective state

We next investigated the affective state of male C57/Bl mice (n = 96, aged 6 weeks at start of experiment) managed for 12 weeks in either groups of 3 per cage (n=24 mice) or singly housed (n=24 mice) with half of each population housed in either conventional or IVC caging systems. To increase the power of the study, we repeated the caging system manipulation in single housed mice only (n=12 mice per condition). After 12 weeks of housing all animals were trained and tested in the reward bias assay over 15 sessions (5 days training, 10 sessions high vs low reward associative learning). After testing in the reward bias assay, mice were also tested using a 2% sucrose preference test and novelty suppressed feeding test (22, 23).

Mice in all conditions showed learning over time (F_(5.454, 125.4)_ = 10.88, p < 0.0001, single housed, conventional caging; F_(5.698, 114)_ = 7.276, p < 0.0001, group housed, conventional caging; F_(5.587, 128.5)_ = 4.222, p = 0.0009, single housed, IVC caging;(F_(9, 207)_ = 4.317, p < 0.0001, group housed, IVC caging). All conditions with the exception of group housed, IVC caged mice exhibited higher accuracy for the high reward association, with a main effect of reward value (F_(1,23)_ = 27.49, p < 0.0001, single housed, conventional caging; F_(1,20)_ = 15.73, p = 0.0008, group housed, conventional caging; F_(1,23)_ = 12.26, p = 0.0019, single housed, IVC caging). Group housed, IVC caged mice showed only a trend level effect (F_(1,23)_ = 4.028, p = 0.0566). Only singly housed, conventionally caged mice showed a time*reward value interaction (F_(5.720,131.6)_ = 2.425, p = 0.0319) associated with higher accuracy over time for the high reward cue.

Consistent with this, comparison of the session 10 data between groups showed that both the conventional housed populations and the single housed IVC populations exhibited a positive reward bias (conventional single t_(23)_=6.947, p<0.001, conventional group t_(23)_=2.276, p = 0.0325, IVC single t_(19)_=2.854, p=0.01 one-sample t-test vs 0% bias) but there was no bias observed for the group housed IVC population. Singly housed, conventional animals had a higher bias score than conventional group housed animals (t_(88)_ = 2.391, p = 0.0189) and single housed IVC animals (t_(88)_ = 2.356, p = 0.0207). There was a main effect of cage type (F_(1,87)_ = 6.491, p = 0.0126) and grouping (F_(1,87)_ = 10, p = 0.0022, single versus grouped conventionally caged, p = 0.0387). There was additionally a trend towards an increase in reward bias in singly housed mice in IVC caging compared to group housed mice in IVC caging (p = 0.0777) and in singly housed, conventionally caged mice compared to singly housed, IVC caging (p = 0.0896). There was no difference between the group housed IVC and conventional cages, but this may have been because both groups had low bias scores. Results from the sucrose preference test were consistent with the reward bias results, showing that singly housed, conventional caged mice had greater reward sensitivity than group housed, conventionally caged mice. There was a main effect of grouping (F_(1,84)_ = 13.83, p = 0.0004, singly housed versus grouped housed conventional animals, p = 0.0015). There was no effect of cage type (p > 0.05). There were no differences in anxiety-like behaviour observed between groups measured by the novelty supressed feeding test. However, there was a trend towards a main effect of cage type on latency to eat (F_(1,81)_ = 3.389, p = 0.0693). Taken together, these data suggest that singly housed, conventionally caged animals are in a more positive affective state than those housed in the other conditions.

In the replication study using single housing in conventional versus IVC housing, both populations showed learning over time and greater high reward accuracy. In singly housed, conventional caged mice, there was a main effect of time (F_(9,135)_ = 12.01, p < 0.0001), reward value (F_(1,15)_ = 7.783, p = 0.0137) and a time*reward value interaction (F_(9,135)_ = 2.857, p = 0.0041, Fig 3A). In singly housed, IVC caged mice, there was a main effect of time (F_(9,135) =_ 9.079, p < 0.0001), reward value (F_(1,15)_ = 5.771, p = 0.0297) but no time*reward interaction (p > 0.05). In line with the above findings, single housing in conventional caging resulted in a greater % reward bias on session 10 compared to single housed, IVC caged mice (t_(30)_ = 2.076, p = 0.0465). IVC caging was also associated with a reduction in % sucrose preference (t_(29)_ = 2.112, p = 0.0434) and higher latency to eat in the novelty supressed feeding test (t_(28) =_ 2.239, p = 0.0333) indicative of reduced reward sensitivity and enhanced anxiety-like behaviour. These findings confirm the previous result suggesting single housed male mice are in a more positive affective state when housed in conventional caging. It remains to be further investigated whether the impacts of IVCs is related to the reduction in social cues e.g. olfactory or auditory, or, a consequence of the environment of the IVC e.g. higher ventilation rates and temperature.

**Fig. 2.**
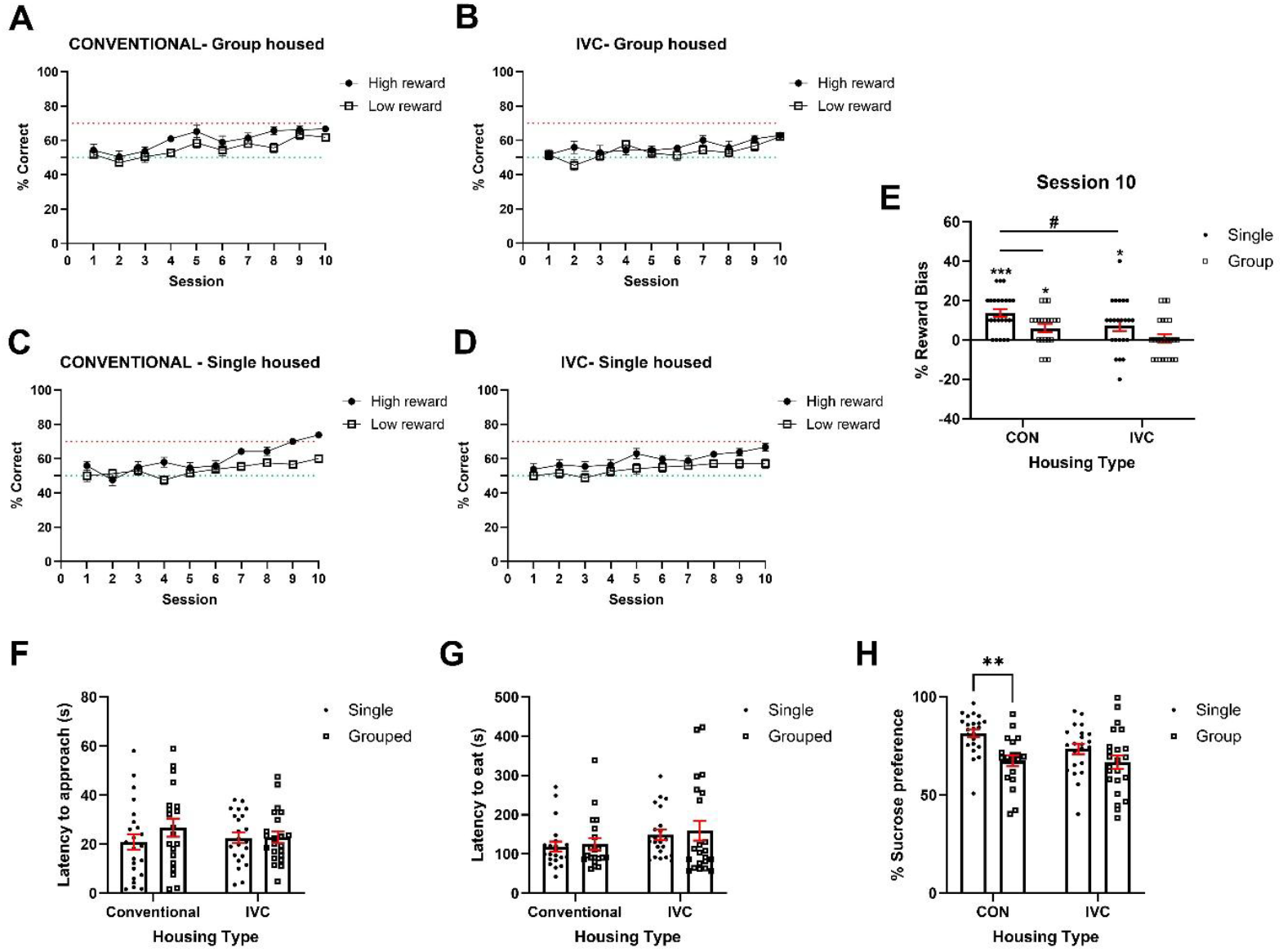
Group housing impaired reward learning and reduced reward sensitivity compared to single housed male mice with individual conventional housing associated with a more positive affective state. Mice were group or singly housed in either conventional or IVC caging for a period of 12 weeks. They were then tested in the reward bias assay, sucrose preference test and novelty supressed feeding test. **A-D** shows learning accuracy across conditions for the high versus low reward over 10 sessions in the four different housing conditions. Only the single housed animals showed a clear separation in the learning rate for the high vs low reward and a reward bias. **E** Group housed mice in conventional caging showed a reduced % reward bias on session 10 compared to singly housed mice in conventional caging. There was no effect of group size or caging on **A** latency to approach or **B** latency to eat from the centrally placed food bowl in the novelty supressed feeding test. **C** Group housing in conventional caging reduced % sucrose preference compared to single housing in conventional caging. *p< 0.05, **p< 0.01 one-sample t-test vs 0% reward bias, #p<0.05 pairwise comparison as indicated. Bars are mean ± SEM with data points overlaid (E, F, G, H).

**Fig.3.**
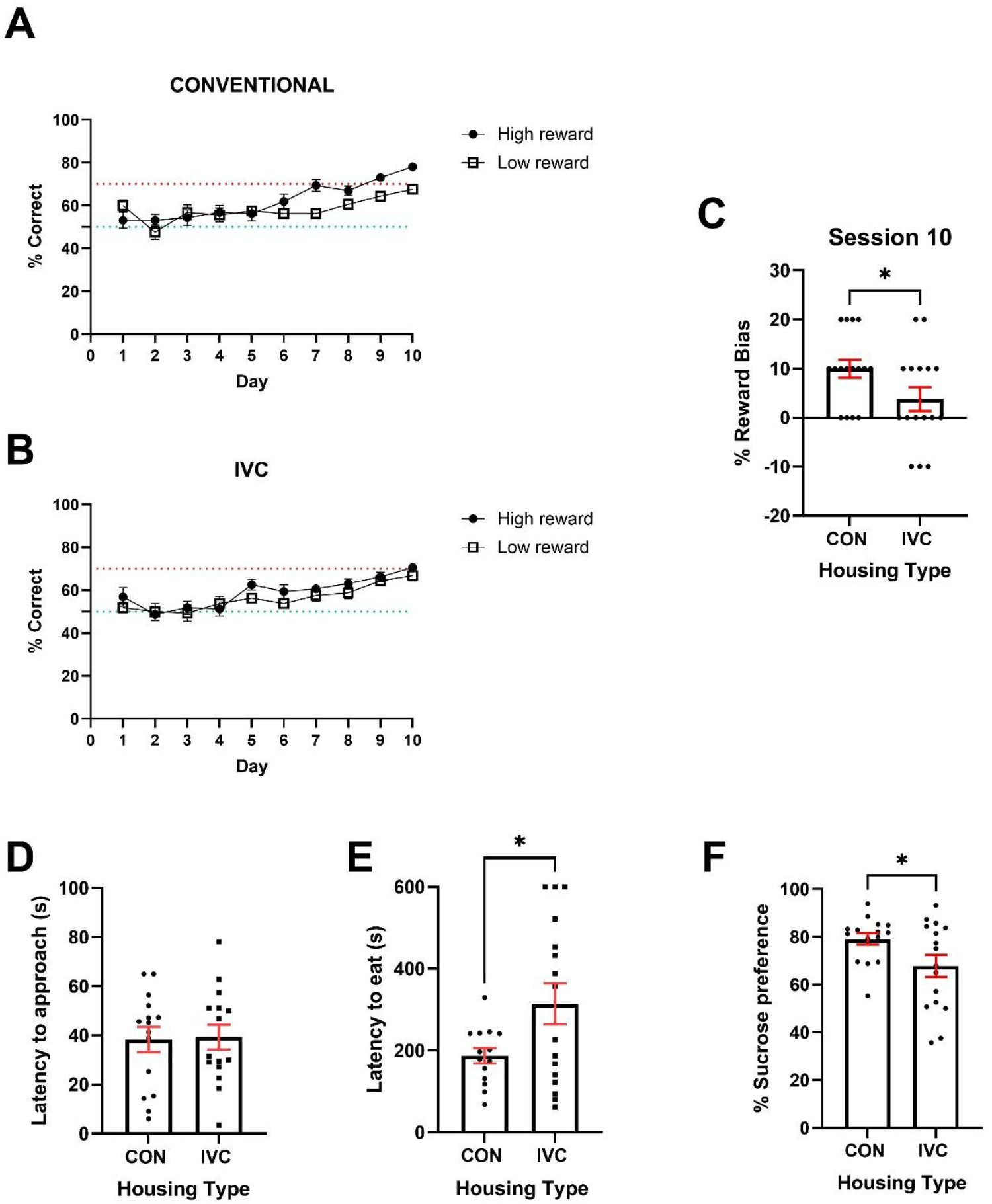
IVC caging impairs reward learning, increases latency to eat in the novelty supressed feeding test and reduces reward sensitivity in single housed male mice compared to conventional caging. Mice were grouped or singly housed in either conventional or IVC caging for a period of 12 weeks. **A&B** shows learning accuracy across conditions for the high versus low reward over 10 sessions. **C** IVC caging reduced % reward bias compared to conventional caging (p<0.05). **D** There was no effect of caging on latency to approach the centrally placed bowl in the novelty supressed feeding test. **E** IVC housed mice had a greater latency to eat from the bowl in the novelty supressed feeding test. **F** IVC caged mice showed a lower % sucrose preference than conventionally housed mice (p < 0.05). *P < 0.05 pairwise comparison. Bars are mean ± SEM with data points overlaid.

## Effects of group vs single housing on ultrasonic vocalisations in male mice

Relatively little is known about communication in mice, but studies have shown they produce a wide range of calls with most emitted in the ultrasonic range (24). In the wild, male mice are thought to use USVs within their territorial behaviours and during courtship (25). Although not an established indicator of wellbeing, we undertook an exploratory study (n=3 cages 3 per cage and n = 3 cages single housed) to investigate if there were differences in the number of USVs emitted by mice housed in groups or individually. We also wanted to see if this changed over time. Recordings were made using a time sampling method with a microphone placed above each cage and data collected for 10min every hour at four timepoints every day. The audiograms were analysed using DeepSqueak and numbers of calls (group housed total calls divided by 3 to account for the number of animals in the cage) plotted against time. There was a main effect of time (F_(1.389,5.556)_ = 14.52, p=0.0081) with evidence that individually housed animals increased their calls over time (time x housing interaction F_(35,140)_ = 6.356, p<0.0001, Fig. 4). Comparing the total number of calls recorded over the duration of the study showed single-housed mice vocalised significantly more than group-housed mice (Mann-Whitney test, *U = 0*, p≤0.05, see Suppl. Fig. S2). Considering the reward bias data these data suggest higher numbers of calls reflect a more positive affective state and monitoring calls could be a useful way to better understand welfare costs of different management approaches as proposed for rats (26).

**Fig. 4.**
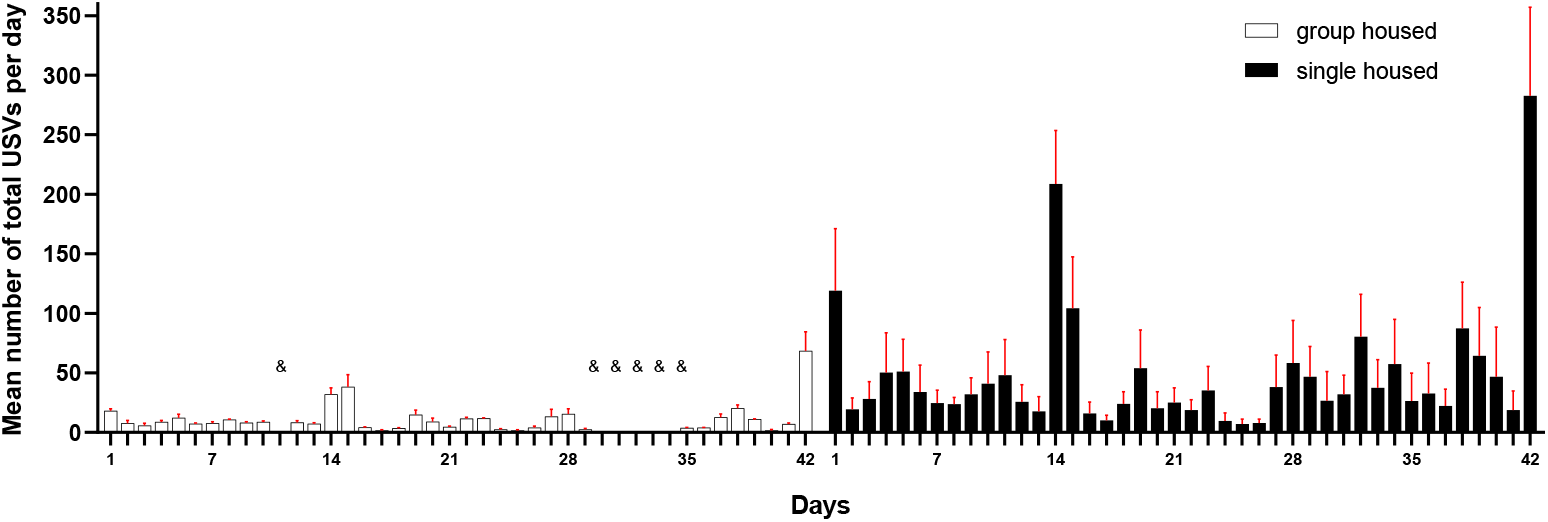
Single housed male mice produce a higher number of calls than group housed male mice over time. Graph illustrates the average number of USV calls per day (4 × 10min recordings) over 42 days of recording. Each bar demonstrates an average from N=3 (white bars: group-housed mice, black bar: single-housed mice). & Missing data due to technical error during USVs recordings.

## Implications for the management of sexually mature male mice

These data suggest male mice do not benefit from social housing when sexually mature and group housed animals experience chronic social stress leading to a depression-like affective state and poorer welfare. In his studies of wild mice, Peter Crowcroft (27) observed that dominant male mice establish and defend territories preventing any other males from being in close proximity. Whilst subordinate animals are tolerated, they avoid physical contact and adapt their circadian behaviours to avoid the dominant male (28). Laboratory mice live within very artificial environments necessitated by both practical and scientific limitations. However, the laboratory environment exposes animals to situations which would not normally be encountered such as an inability to express normal escape or subordinate behaviours and stress due to human interventions which can lead to high levels of aggression (29, 30).

The consensus in the current literature and most country’s guidance and ethical regulation is that mice should not be housed individually unless they are a particularly aggressive strain or for specific scientific objectives (1-3). This is based on the premise that individually housed mice will be in a poorer welfare state versus group housed mice even with some inter-male aggression. The rationale for this is a body of literature which suggests that male mouse behaviour and physiology is changed when they are individually housed, and these reflect detrimental effects on the animal’s welfare (4, 31). However, much of this work is not based on objective assessments of affective state or stress but rather indirect measures of different scientific endpoints or social interaction tests. Studies where social interaction, choice paradigms or experiments involve motivation to obtain access to social contact may also be more complex than might initially be apparent. For example, wild male mice are highly territorial and engage in behaviours to defend their territory and will approach any animal which enters the area (27) In laboratory mice, these socially motivated tasks may reflect social interactions associated with territorial behaviour as opposed to seeking positive social contact (29, 30). Van Loo (31) reports on high levels of sniffing at the openings in and alongside partitions between mice and digging at sawdust which they describe as ‘attempts to make social contact’ however, this behaviour could equally be driven by aggression confounding interpretation of the data.

## Conclusions

These studies provide an objective assessment of the welfare costs of group versus single housing of male C57/Bl mice with evidence that a more positive affective state is observed when animals are singly housed. These findings argue against established ideas about the sociability of male mice and the welfare benefits of social housing. This has important implications in terms of colony management as well as impacting scientific outcomes and the translational validity of studies in mouse models managed following current guidelines. Our findings also show that IVC caging systems are associated with poorer welfare in both single and group housed male mice. It is important that the welfare costs of these management systems are now reconsidered in line with the principles of the 3Rs.

## Methods

### Complex Cue T-Maze (CCTM)

To quantify affective state in mice, a novel behavioural test was developed in-house to assess reward learning and memory over time. The complex cue T-Maze apparatus (CCTM, patent pending, fig 1) was developed to test the mouse’s ability to pair a tactile stimulus (3D patterned floor) with the delivery of a high versus low value liquid reward. Mice were first habituated to the arena during a 10 min session of free exploration. They then began the training phase.

### Training Phase

To enhance engagement with the task, mice were placed on a mild food restriction regime. Mice received 2.5g standard lab chow (Purina, UK) per day 1 hour post testing. Mice were given *ad libitum food* on non-testing days (across the weekend). Mice underwent 5 sessions of basic discrimination. Here, they were presented with a smooth blank floor (no 3-D pattern) in one arm versus a patterned floor in the other. At the start of the test, the mouse is placed in the start box with all doors closed. The trial begins when the door to the goal arms was manually opened. Once the mouse entered one goal arm (all four feet), access to the other arm is closed. The mouse was rewarded with 10 µl of 10 % condensed milk for choosing the patterned floor. Condensed milk was manually dispensed into the arm via a reward port using a Gilson pipette. No reward was given for choosing the blank floor, however the pipette was still raised to the port to prevent formation of association of experimenter movement with reward delivery.

The correct floor position (left or right) was presented in a pseudorandom order for 20 trials with a repeated trial following an incorrect response. If the mouse chose the pattered floor 70% or more of the time by the 5^th^ day they were considered to have successfully discriminated between floor and proceeded to reward learning bias testing.

### Reward Learning Phase

Mice underwent 10 sessions where they were presented with 4 different 3-D pattered floors (Suppl. Fig. S1.). These floors were split into two pairs: a high reward (10ul, 5% condensed milk) pair and a low reward (10ul, 2% condensed milk) pair. For each pair, one floor was rewarded (CS+) and the other was unrewarded (CS-).

20 trials per session were undertaken with both pairs of floors presented within the same session (10 times per pair). Presentation of the pairs, as well as position of the rewarded floor (left or right) were pseudorandom. There was no repeat trial for incorrect response. The number of times the mouse correctly chose the high or low rewarded floor versus their respective blanks was expressed as ‘% correct’. This was calculated by (number of correct choices/total number of trials) x 100. A reward bias was calculated by subtracting % correct for high reward versus % correct for low reward. Reward bias was calculated for the final testing session. N = 4 (n = 2 per group) mice from the singly housed mice in IVC vs conventional caging study were excluded from the study due to inability to learn basic discrimination. These mice were excluded from all behavioural measures.

### Novelty Supressed Feeding Test

The testing apparatus consisted of a white Perspex^®^ arena (60 cm ×60 cm ×30 cm). The floor was covered with approximately 1 cm of sawdust bedding. Twenty three hours prior to behavioural testing, all food was removed from the home cage. At the time of testing, two pellets of food (regular chow) were placed on white filter paper positioned in the centre of the box. Subject mice were placed in a corner of the box, and the times elapsed before approaching and eating the food were measured. Approaching was defined as the nose being within 1cm of the food pellet. Mice were removed from the arena once eating had commenced or after a maximum of period of 10 minutes.

### Sucrose Preference Test

The test consisted of three consecutive habituation days and on the fourth day, a preference test session. Access to water was removed from the mice the evening before experiment to encourage water consumption. Food was provided ad libitum. On the first day, mice were removed from their home cage and placed in a fresh cage, containing sawdust and a cardboard tube, with sipper sacks (450ml, Edstrom Industries Inc) and metal nozzles (2cm) placed to the left and right of the cage, containing 2% sucrose solution made up in tap water. Mice were left for 4h, and water intake was monitored to ensure consumption from both sides. At the end of the 4h session mice were returned to their home cage. This process was repeated on the second day, but with plain tap water. This was switched back to 2% sucrose on the third day. Sawdust was shaken between days.

On the preference test day, both tap water and 2 % sucrose solution were available. Position of sipper sacks was counterbalanced. At 1h and 2h the position of the sipper sacks was switched. Water intake was measured at 1h, 2h and 4h. % sucrose preference was calculated using: (total amount of sucrose solution consumed/total amount of liquid consumed) *100.

### Experiment 1: Validation of the complex cue T maze reward learning assay Subjects

Male C57BL/6 mice were purchased from Envigo, UK at 8 weeks old. 18 mice were tested (n = 9 treatment group, n = 9 control group). Mice were individually housed in conventional caging under a 12 h reverse light/dark cycle and had access to food and water *ad libitum* with the exception of during experimental testing as described above. Enrichment was consistent with the above. Testing took place during the dark phase between 8am and 5pm.

### Corticosterone treatment and CCTM testing

Animals underwent 5 days of basic discrimination in the CCTM as described above. Mice were randomly assigned to the treatment or control group and the experimenter was blind to treatment. The treatment group were given 25ug/ml corticosterone-HBC (C174, Sigma) in their drinking water. The control group were given tap water only. Water was provided in dark drinking bottles to prevent exposure of corticosterone to the light. Water was changed every 2 days. Treatment began 14 days before onset of behavioural testing. Following this, mice underwent 10 days of reward learning bias testing in the CCTM as described above. Treatment continued throughout behavioural testing. N = 3 mice per treatment did not complete the 10^th^ day of behaviour due to technical error.

### Experiment 2: Assessment of the impact of group size and caging type on reward learning, novelty suppressed feeding, and sucrose preference as indicators of affective state

#### Subjects

Male C57BL/6 mice were purchased from Envigo, UK at 6 weeks old. 8 replicates were tested (total n= 96) with each cohort comprising n = 12. Upon arrival, mice were evenly and randomly assigned to one of four groups: individually housed in conventional caging, individually housed in IVCs, group housed (3 per cage) in conventional caging or group housed in IVCs. All mice were kept in the same room and were provided with environmental enrichment consisting of nesting material (50% paper bedding (IPS, UK) and 50% Sizzlenest bedding (Datesand, UK), two cardboard tubes suspended from the cage lid and a chew block. Mice were housed in these conditions for 9 weeks total (see behavioural timeline, fig.5). Mice were housed under a 12 h reverse light/dark cycle and had access to food and water *ad libitum* with the exception of experimental testing. Testing took place during the dark phase between 8am and 5pm. Where fighting occurred and led to significant injury, mice were separated upon advice from the Named Veterinary Surgeon (NVS) and excluded from the study (n = 3, S1).

**Fig. 5.**
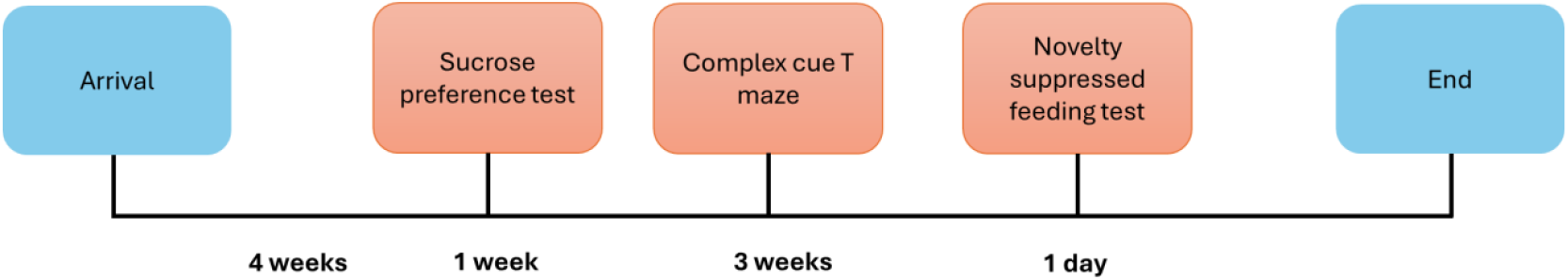
Behavioural timeline for Experiment 1.

To further investigate the observed differences between singly housed mice in IVC vs conventional caging, a further 36 mice were tested (18 per group). They were housed for 9 weeks with the same enrichment as described above. All procedures were carried out under local institutional guidelines (approved by the University of Bristol Animal Welfare and Ethical Review Board) and in accordance with the UK Animals (Scientific Procedures) Act 1986.

### Experiment 3: Ultrasonic vocalisation study

#### Subjects

12 male C57BL/6JOlaHsd mice (Envigo (UK) from the age of 6 weeks till 12 weeks were used in the USVs experiment. Mice were managed under 12hr:12hr reverse lighting (lights on 8:15) with *ad libitum* food and water and basic enrichment (nesting material, cardboard house, wood chew block, cardboard tube). The USV study used a sample size of 3 cages per group with single-housed mice (N=3) and group housed mice 3 animals per cage (N=3). All behavioural studies were carried out during the animal’s active phase. All experiments were performed in accordance with the Animals Scientific Procedures Act (ASPA, UK) 1986 and were approved by the University of Bristol Animal Welfare and Ethical Review Body and UK Home Office (PPL number PP4130557).

#### USVs recording and analysis

Ultrasonic vocalisations were recorded over 42 days from home cages four times a day at 9:00, 12:00, 15:00, 18:00 for 10 minutes each by using 6 high-frequency microphones (2- to 200-kHz range, CM16/CMPA UltraSoundGate Condenser Microphone, Avisoft Bioacoustics, Germany) suspended 3cm above the cage lid to cover recordings from all home cage space. Microphones were connected to computers via ultrasound recording interfaces (416H UltraSoundGate, Avisoft Bioacoustics, Germany). The acoustic data was recorded by Avisoft RECORDER USGH software (version 4.4, Avisoft Bioacoustics, Germany) and batch analysed by generating 290 spectrograms in Deep Squeak software (version 3.6.2.). The analysis parameters based on previously established literature (32) were set as follows: a total analysis length of 0, an analysis chunk length of 6, a frame overlap of .0001 s, a frequency low cut-off of 20 kHz, a frequency high cut off of 110 kHz, and a score threshold of 0. Once the spectrograms were generated, the files were processed by rejecting false calls and background noise of tonality <0.25. Then spectrograms were manually processed, where a trained experimenter individually inspected each .wav file to remove the remaining background noise. If a detected vocalization was not fully contained by the automatic detection boxes, then the detection boundaries would be redrawn in order to encompass the entirety of the USV. For the USVs analysis, the experimenter was blinded to the housing groups.

### Statistical analysis

Data were checked for normality using the Shapiro-Wilk test. If more than half of experimental groups were non-normally distributed, a non-parametric test was applied. Where data were normally distributed and required single factor analysis between 2 groups, an unpaired t-test was used. Where two-factor analysis was required (group size*cage type, reward value*time, housing*time) a two-way ANOVA or repeated measures two-way ANOVA was used. If data points were missing due to a random event, a mixed-effects ANOVA was carried out instead. Where significant main effects occurred (p < 0.05), Šídák’s multiple comparisons test were carried out. Trend level effects (p < 0.1) are reported but not further analysed. Reward bias scores from the final session (session 10) were also analysed using a one-sample t-test versus a theoretical mean of 0% bias to determine whether animals developed a reward-induced bias. Where data was not normally distributed a Mann-Whitney U test was performed. Outliers were defined as data points ± 2SDs away from the mean and were excluded from analysis. All statistical exclusions are outlined in **S1**. The statistical unit refers to the individual mouse. Data were graphed and analysed using GraphPad Prism v 10.0.03.

## Acknowledgments

This research was supported by funding from the NC3Rs reference NC/Y00082X/1

## Conflict of interest statement

ER, MJ, JD, JH are creators of the 3Hs initiative, a framework designed to promote refined housing, handling and habituation methods. This initiative includes working with commercial suppliers of products used in the management of laboratory animals. ER has received funding for collaborative and contract research from pharmaceutical companies, Boehringer Ingelheim, Compass Pathways, Eli Lilly, IRLabs Therapeutics, Pfizer and SmallPharma and acted as a paid consultant for Compass Pathways and Pangea Botanicals.

## Supplementary material

Video 1: An example of a mouse running a trial in the CCTM

## Supplementary figures

**Supplementary Fig. S1.**
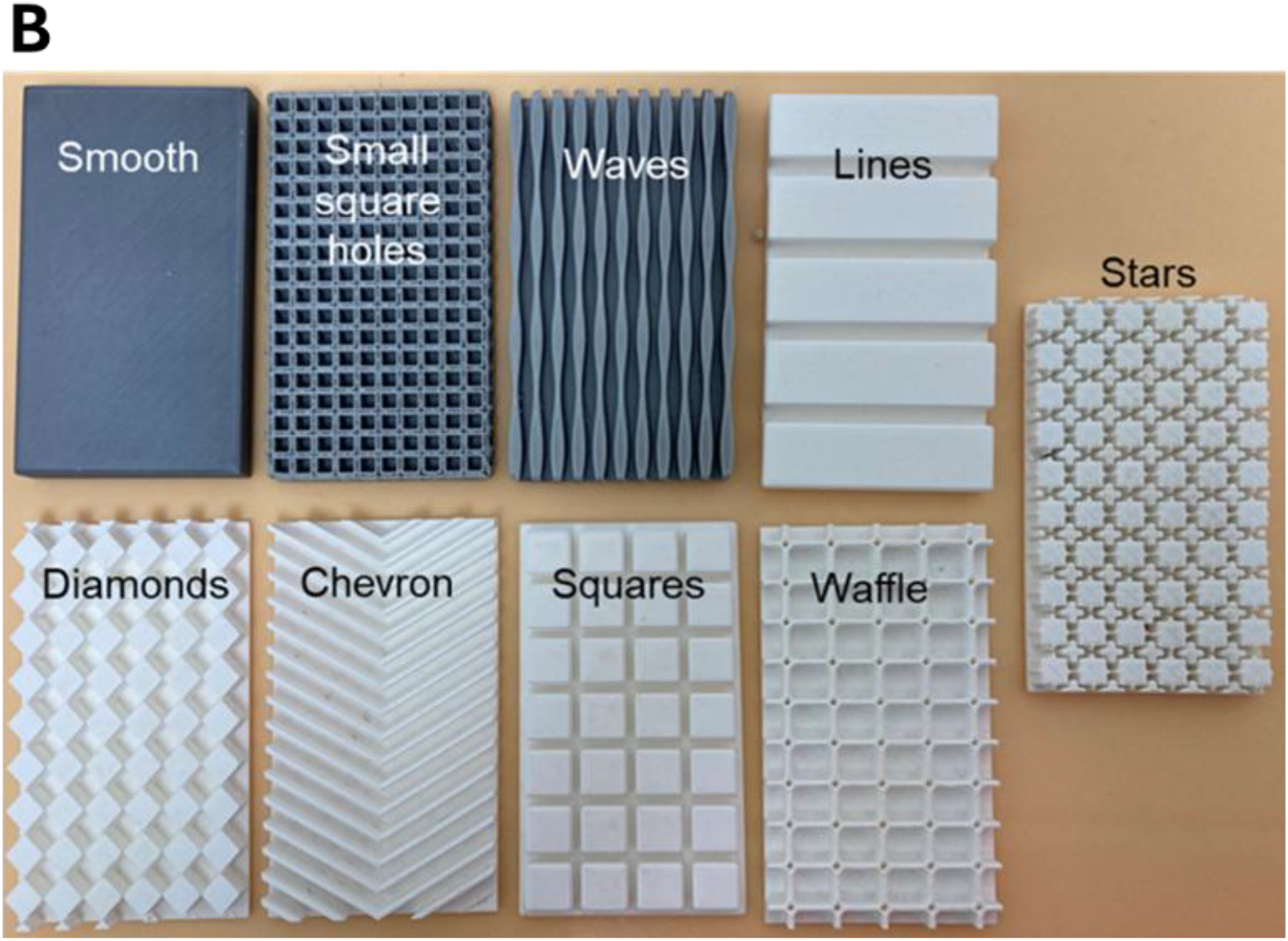
Complex Cue T Maze 3D printed floors, dimensions =. 13 cm x 7.5 cm x 1 cm.

**Supplementary Fig. S2:**
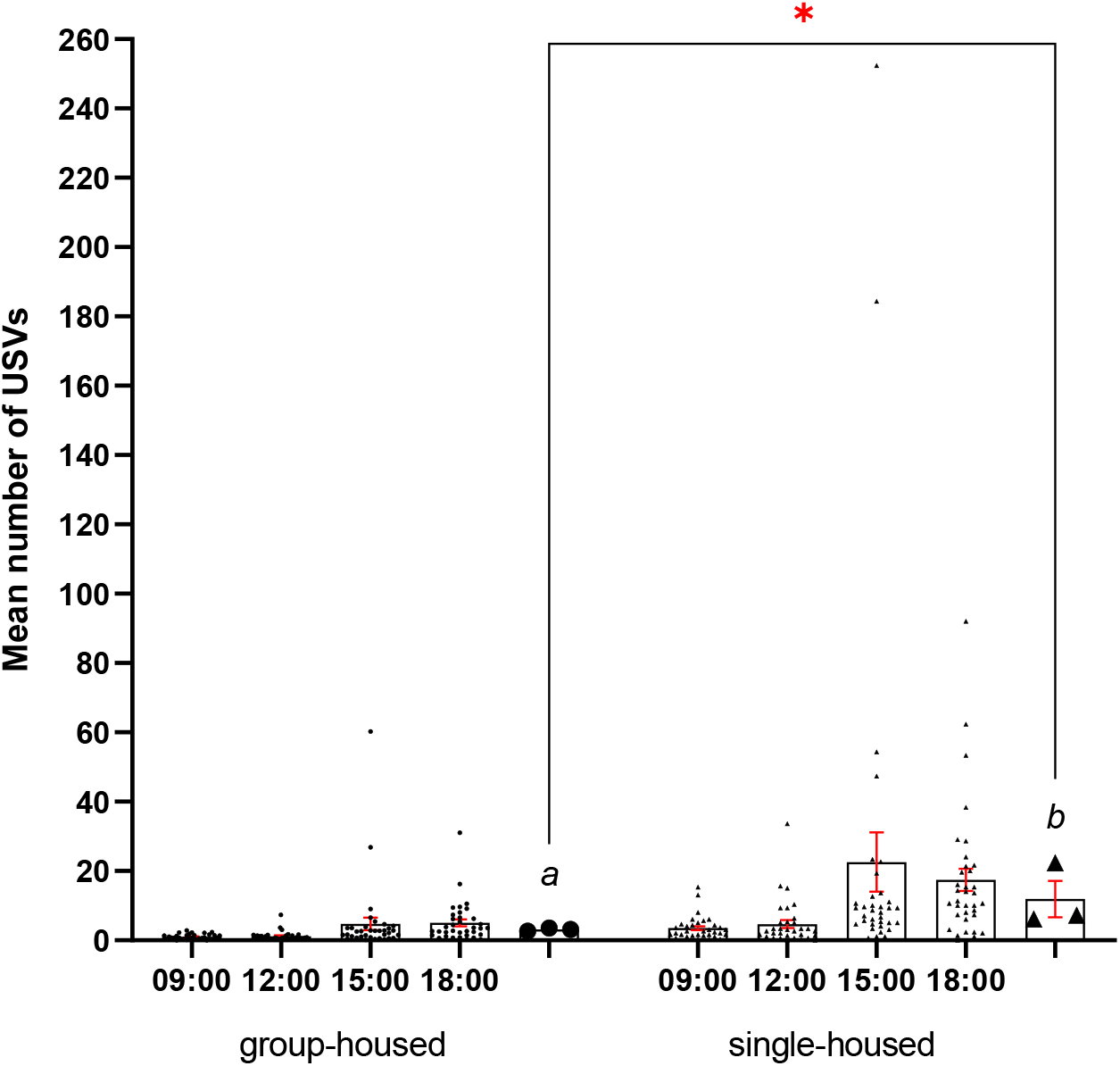
Single-housed mice vocalised more than group-housed mice. The graph illustrates mean number of calls for each timepoint (9:00, 12:00, 15:00, 18:00) per housing group (both N=3) over 42 days of recording. Additional bars a and b represent the mean number of calls per day (average of 4 timepoints) per all recording period. Data from group-housed mice was normalised to N=1, by dividing the total number of calls by 3. The data for each time point is represented by individual data points (circle - group-housed mice, triangle - single-housed mice) and the mean bars with standard error of the mean. *p≤0.05

**Supplementary Table S1.**

| Experiment | Test/measure | Exclusion |
| --- | --- | --- |
| Housing and grouping comparison | Latency to sniff - NSFT | Outliers: CON, S – n=1, IVC, S – n=2, CON, G – n=1, IVC, G – n=2<br>Pre-exclusion: CON, G n=3 |
|  | Latency to eat - NSFT | Outliers: CON, S – n=3, IVC, S – n=2, CON, G – n=1, IVC, G – n=2<br>Pre-exclusion: CON, G n=3 |
|  | % preference - SPT | Outliers: CON, S – n=1, IVC, S – n=2, CON, G – n=1, IVC, G – n=1 |
|  |  | Pre-exclusion: CON, G n=3 |
|  | Reward bias | Outliers: CON, S – n=0,<br>IVC, S – n=1, CON, G – n=1,<br>IVC, G – n=0<br>Pre-exclusion: CON, G n=3 |
| IVC vs Conventional direct<br>comparison | Latency to sniff - NSFT | CON – n=1, IVC – n=1 |
|  | Latency to eat - NSFT | CON – n=2, IVC – n=0 |
|  | % preference - SPT | CON – n=3, IVC – n=2 |
|  | Reward bias | None |
| CORT (+/-) | Reward bias | None |

**S1. Statistical exclusions per experiment and output measure.** CON-conventional, IVC-individually ventilated caging, NSFT-novelty suppressed feeding test, SPT-sucrose preference test.

